# Temporal Prediction through Integration of Probability Distributions of Event Timings at Multiple Levels

**DOI:** 10.1101/2024.10.15.618601

**Authors:** Yiyuan Teresa Huang, Zenas C. Chao

**Affiliations:** International Research Center for Neurointelligence (WPI-IRCN), UTIAS, The University of Tokyo, 113-0033, Japan; Department of Multidisciplinary Sciences, Graduate School of Arts and Sciences, The University of Tokyo, Tokyo, Japan; Japan Society for the Promotion of Science (JSPS), Japan

## Abstract

Our brain uses prior experience to anticipate the timing of upcoming events. This dynamical process can be modeled using a hazard function derived from the probability distribution of event timings. However, the contexts of an event can lead to various probability distributions for the same event, and it remains unclear how the brain integrates these distributions into a coherent temporal prediction. In this study, we create a local-global foreperiod paradigm consisting of a sequence of paired trials, where in each trial, participants respond to a target signal after a specified time interval (i.e. foreperiod) following a warning cue. The prediction of the target onset in the second trial can be based on the probability distribution of the second foreperiod (local level) and its conditional probability given the foreperiod in the first trial (global level). These probability distributions are then transformed into hazard functions to represent the local and global temporal predictions. Reaction times to the target signal are best explained by incorporating both local and global predictions, indicating that both levels of temporal information contribute to making predictions. We further show that electroencephalographic source signals are best reconstructed when integrating both predictions. Specifically, the local and global predictions are separately encoded in the posterior and anterior brain regions, and to achieve synergy between both predictions, a third region—particularly the right posterior cingulate area—is needed. Our study reveals brain networks that integrate multilevel temporal information, providing a comprehensive view of hierarchical predictive coding of time.

## Introduction

Predicting when an event will occur is a fundamental cognitive process that allow for the efficient allocation of attentional resources and optimal motor performance (Morillon & Schroeder, 2015; Sørensen et al., 2015; Vangkilde et al., 2012). It has also been found that precise temporal predictions optimize the processing of sensory information by enhancing contrast sensitivity (Rohenkohl et al., 2012). To study temporal prediction, researchers often use a foreperiod task where participants are required to press a button as soon as a target signal appears following a warning signal. The time interval between the warning and the target signals is referred to as the foreperiod. To achieve a fast response, it is crucial to learn the probability distribution of the foreperiod as participants can use it to expect the target onset. In this task, temporal prediction is often modeled by a hazard function (HF), which represents the ongoing updates to predictions as time progresses (Luce, 1991; Nobre et al., 2007). This function, derived from the probability distribution of the foreperiod, represents the conditional probability that the target signal will occur, given that it has not yet happened.

For neural correlates of the temporal prediction, higher HF values (i.e., stronger prediction) have been associated with increased activity in the parietal area, as observed in single-neuron recordings and human MRI studies (Coull et al., 2016; Janssen & Shadlen, 2005). Additionally, increased alpha power in human EEG has also been linked to higher HFs (Herbst & Obleser, 2017). Moreover, as temporal predictions elapse over time, these dynamics can be tracked in the brain by training time-solved EEG signals in a forward encoding model. This modeling approach can effectively distinguish brain signals associated with varying HFs derived from different probability distributions (Herbst et al., 2018). However, the timing of an event may be influenced by multiple probability distributions, particularly when considering the specific contexts involved. Take Beethoven’s Symphony No. 5 as an example. It begins with a “short-short-short-long” motif, commonly known as “fate knocking at the door.” To predict the fourth interval, one might expect a higher chance of “short” when only considering the probability distribution of each element, but there could be a higher chance of “long” when considering the conditional probability of the multi-element pattern that characterizes the motif. It raises a question of how the brain integrates various probability distributions for the same event across different levels, to generate a coherent temporal prediction.

To understand how the temporal prediction is established by probability distributions of event timings across multiple levels, we introduce a local-global foreperiod paradigm. In this paradigm, two foreperiod trials (FP1 and FP2) are paired in a sequence, leading to two levels of statistical regularities. Predicting FP2 can be based on not only the probability distribution of the single foreperiod (local level), but also the probability distribution of FP2 conditioned on the preceding foreperiod (i.e., FP1) when the sequence structure is considered (global level). We obtain two hazard functions from the two probability distributions to represent local and global temporal predictions, respectively, and then model reaction times to the target and EEG source signals. Using a model-fitting approach, both behavioral and neural results indicate that the two statistics are learned jointly, rather than independently, for prediction establishment. Furthermore, we find that local and global temporal predictions are encoded in distinct brain regions, with additional brain regions processing their interaction. Our study offers an experimental platform for exploring multilevel temporal prediction, and identifies key brain regions involved in the hierarchical predictive coding of time.

## Results

### Local-global foreperiod paradigm to establish multilevel temporal predictions

To control predictability of an upcoming event based on two probability distributions calculated at different levels, we created a novel local-global foreperiod paradigm. During each trial, participants received a warning signal (a low-pitched tone and a white dot on the screen) and were instructed to press the button promptly as the target signal (a high-pitched tone and a red dot) appeared. The delay between the warning and the target signal is referred to as the foreperiod (Figure 1A). Two consecutive trials are considered a “sequence” when the warning signal from the second trial occurs 1.2 seconds after the target signal from the first trial, with the interval between two consecutive sequences ranging from 3 to 3.2 seconds (Figure 1B, top). Within each sequence, the foreperiods for the first and the second trials are denoted as FP1 and FP2, respectively. For each condition, a block of 50 sequences (100 trials) were delivered.

**Figure 1.**
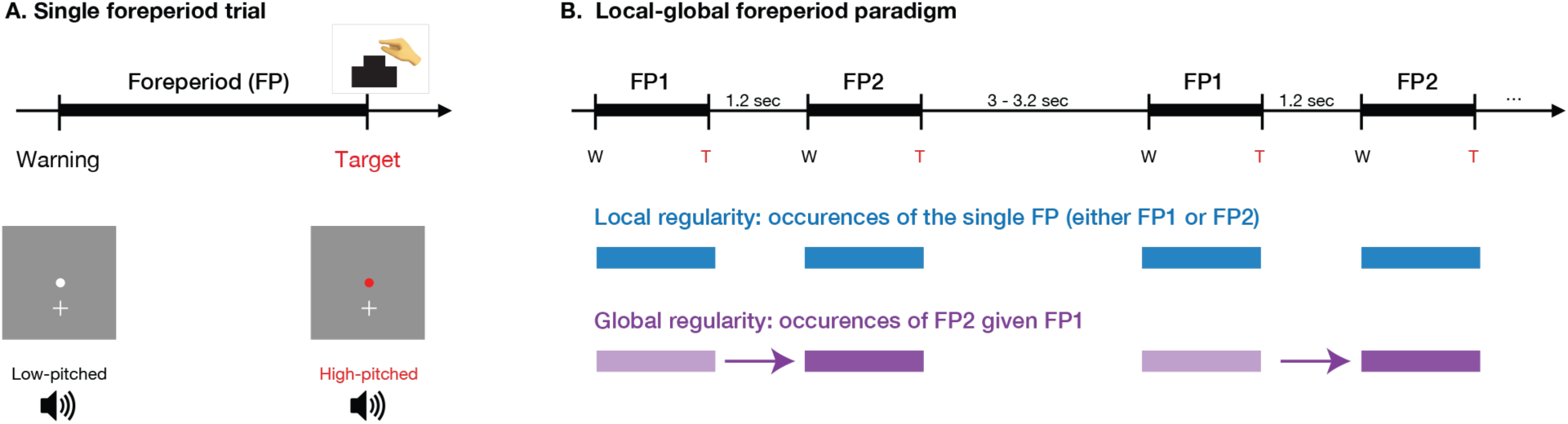
Task design. (A) a single foreperiod (FP) trial consisted of a white dot and a low-pitched tone as a warning signal, and a red dot and a high-pitched tone as a target signal. The FP is the time interval between the warning and the target, ranging from 0.4 to 2 sec. Participants were requested to press a *zero* button when the target appeared. (B) A sequence consisted of two FP trials (denoted as FP1 and FP2) separated by a short blank, and a longer blank separated two sequences. For the local regularity, regardless of the sequence structure, the probability distribution was computed based on single FP occurrences (either FP1 and FP2) (highlighted with the blue rectangle). For the global regularity, the probability distribution was computed based on FP2 occurrences (the purple rectangle) given on long or short durations of FP1 (the light purple rectangle).

This sequence design allows the predictability of FP2 to be established with two distinct probability distributions (Figure 1B, bottom). The first is a local-level probability distribution, derived directly from the frequency of the single FP2, regardless of the sequence structure. This is represented by the probability density function PDF_L_. The second is a global-level probability distribution, incorporating sequence structure by calculating the conditional probability of FP2 given FP1, the preceding trial. This is expressed through the conditional probability density function PDF_G_.

Four sequence blocks were created, each with a unique configuration of PDF_L_ and PDF_G_ (Figure 2). This block design allows us to disentangle the effects of PDF_L_ and PDF_G_. Taking Block 1 as an example (Figure 2A), the foreperiods were set either as short (between 0.4 and 1.1 seconds, denoted as *S*) or long (between 1.3 and 2 seconds, denoted as *L*). Two sequence types were used: *LL* (long FP1 and long FP2) and *SS* (short FP1 and short FP2), each presented 25 times. For the local-level probability, there were 25 trials of *S* and 25 trials of *L*, resulting in PDF_L_ of 50% for *S* and 50 % for *L* (denoted as 50-50 PDF_L_; Figure 2B). For the global-level probability, long FP2 always followed long FP1, and thus PDF_G_ for FP2 given long FP1 was a skewed unimodal distribution of almost 100% *L*. Similarly, short FP2 always followed short FP1, and thus PDF_G_ for FP2 given short FP1 was a skewed unimodal distribution of almost 100% *S*. The other three blocks consisted of different numbers of sequences *LL* and *SS*, with additional sequences *LS* and *SL*. While PDF_L_ was always controlled at an even 50-50 ratio, as in Block 1, PDF_G_ varied. Furthermore, to model the temporal dynamics of the local and global predictions, PDF_L_ and PDF_G_ in the four blocks were transformed into local and global hazard functions (denoted as HF_L_ and HF_G_) (Figure 2C; see the formula in Methods). Please note that predictability of FP1 is solely determined by PDF_L_ and HF_L_, which are the same across all blocks.

**Figure 2.**
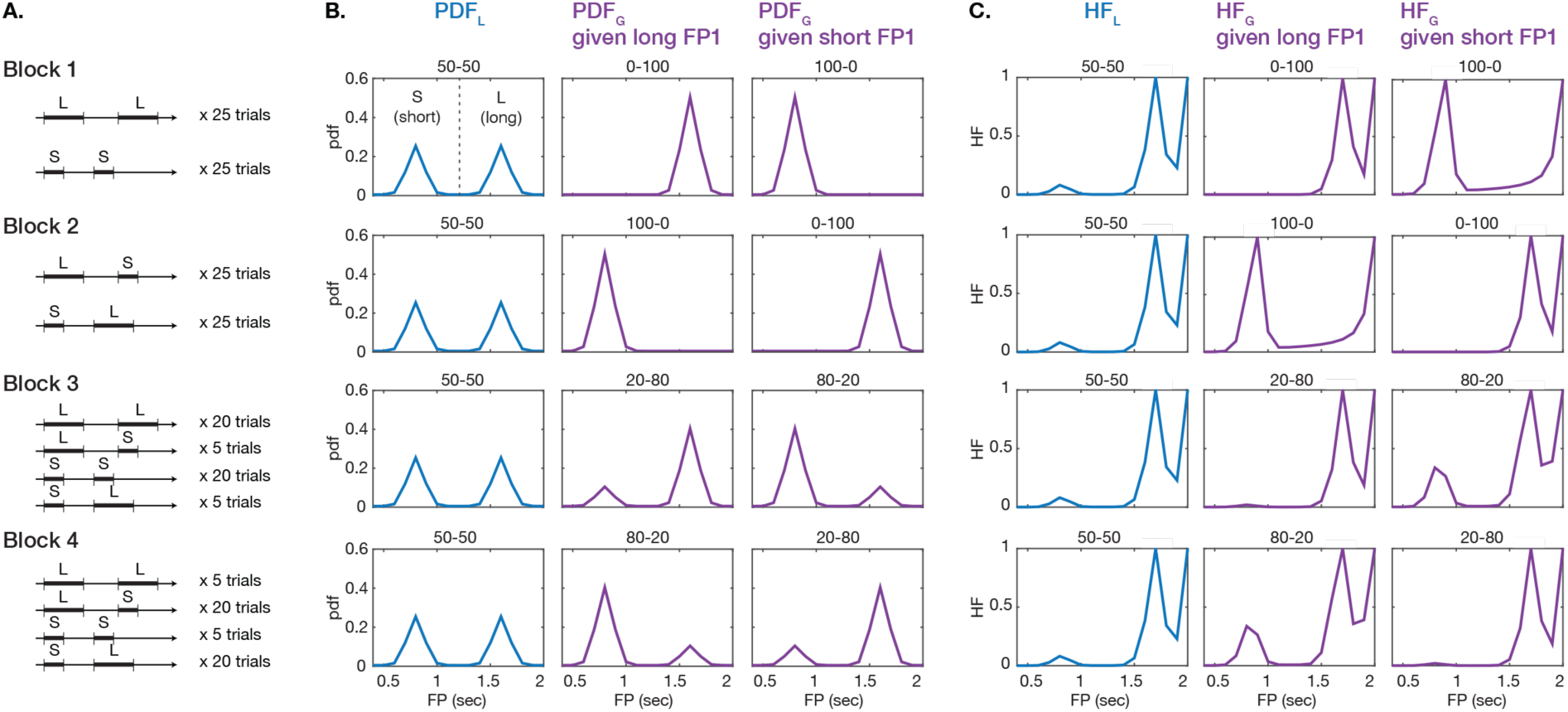
Block design for different probability distributions and hazard functions. (A) Four blocks were created, each containing 50 FP sequences. The FPs timed between 0.4 and 1.1 sec and between 1.3 and 2 sec are denoted as S and L, respectively. There are four types of FP sequences with different trial configurations for each of the four blocks. (B) Probability distributions computed based on the local and global regularities are denoted as PDF_L_ (in blue) and PDF_G_ (in purple), respectively. Two PDF_G_ were calculated, one given long FP1 and the other given short FP1. The number on each plot (e.g., 50-50) describes the proportion of cumulative probabilities in the range of S and L. (C) Hazard functions were computed based on the respective probability distributions in the panel B. HF_L_ and HF_G_ are denoted as computed based on PDF_L_ and PDF_G_, respectively.

Thirty-one participants (16 males and 15 females; age: 23 ± 2.9 years old, mean ± standard deviation) were included in the study. During the experiment, the four blocks were delivered in a random order and repeated three times, each with a different random order, resulting in a total of 12 block representations (run). At the end of each run, participants were asked to identify which sequence types were more frequent to confirm their recognition of the sequence structure. The accuracy was 87 ± 14% (mean ± std, n= 31 participants, chance level= 50%). During the task, we also recorded reaction time to the target signal and 64-channel EEG signals.

### Reaction time explained by both local and global predictions and their integration

To investigate the use of local and global temporal information in predicting FP2, we analyzed the correlations between and the values of HF_L_ and HF_G_ and the reaction times to the target following FP2 (later we simply refer to as reaction times following FP2). Reaction times were first log-transformed to reduce concerns regarding the assumption of a normal distribution (Whelan, 2008). Then, we removed the first 20 trials from each block to account for participant adaptation. Additionally, we excluded any outlier trials where reaction times exceeded 2.5 standard deviations. The average reaction times following FP1 and FP2 are shown in Supplementary Table 1.

Before focusing on FP2, we first verified whether the prediction of FP1 could be explained by local prediction, as studies with a single foreperiod have shown that faster reaction times are associated with higher hazard rates. To achieve this, we used a linear-mixed effect model to regress the reaction times following FP1 against the fixed effect, HF_L_ values, and the individual difference was considered by setting the participant set to the random effect. A significantly negative effect of HF_L_ was found, indicating that faster reaction following FP1 was associated with stronger local prediction (*p* < 0.001, n = 31 participants, see Table 1). This is consistent with results from other studies.

**Table 1.** Effect of HF_L_ on reaction times following FP1.

|  | Estimates | SE | *t value* | *p* | *Adj R*<sup>2</sup> |
| --- | --- | --- | --- | --- | --- |
| (Intercept) | 5.44 | 0.02 | 273.96 | <0.001 | 0.292 |
| HF<sub>L</sub> | -0.08 | 0.01 | -15.93 | <0.001 |  |

*n* = 13641 observations. Random effect: participants. Estimate: correlation coefficient (beta value). SE: standard error. Adj R<sup>2</sup>: adjusted R-squared.

Next, we examined how HF_L_ and HF_G_ contribute to the reaction time following FP2. We used linear-mixed effect models to regress the reaction times following FP2 against 4 different sets of variables: (1) HF_L_ (denoted as HF_L_-only), (2) HF_G_ (denoted as HF_G_-only), (3) HF_L_ and HF_G_ (denoted as HF_L_+HF_G_), and (4) HF_L_ and HF_G_ and their interaction (denoted as HF_L_+HF_G_+HF_L_*HF_G_). Please note that in our regression analysis, the duration of FP1 (*L* or *S*) was used as a covariate. This adjustment was necessary because we observed that the duration of FP1 influenced the response following FP2, which is known as the asymmetrical sequential effect (Los & van den Heuvel, 2001). Specifically, reaction times following a short FP2 were consistently longer after a long FP1 (as in the sequence *LS*) compared to after a short FP1 (as in the sequence *SS*) (see Supplementary Table 2). By including the duration of FP1 as a covariate, we were able to control for this confounding effect. Additionally, the participant was assigned as a random effect for controlling individual differences.

Model comparisons (see Table 2) show that the model that incorporated HF_L_, HF_G_, and their interaction can best predict the reaction time following FP2 (with the smallest AIC and BIC for model fitness and the highest adjusted R squared for model explanation). The results of the best model are shown in Table 3. The interaction term (HF_L_*HF_G_) had a notably positive coefficient (*p* < 0.001), indicating that the effect of HF_L_ on the reaction time is moderated by the value of HF_G_. Specifically, the effect of HF_L_ on the reaction time was weaker under greater HF_G_ values, but stronger under smaller HF_G_ values. In other words, local prediction had a reduced impact on behavioral responses when global prediction was strong. For HF_L_ and HF_G_, their main effects on FP2 were significantly negative (*p* < 0.001), indicating that higher hazard values, either local or global, led to faster responses. These findings suggest that the prediction of FP2 involves both local and global probability distributions. Importantly, local and global predictions interact to influence behavioral responses, suggesting an integration process that enables mutual modulations between the two prediction levels.

**Table 2.**
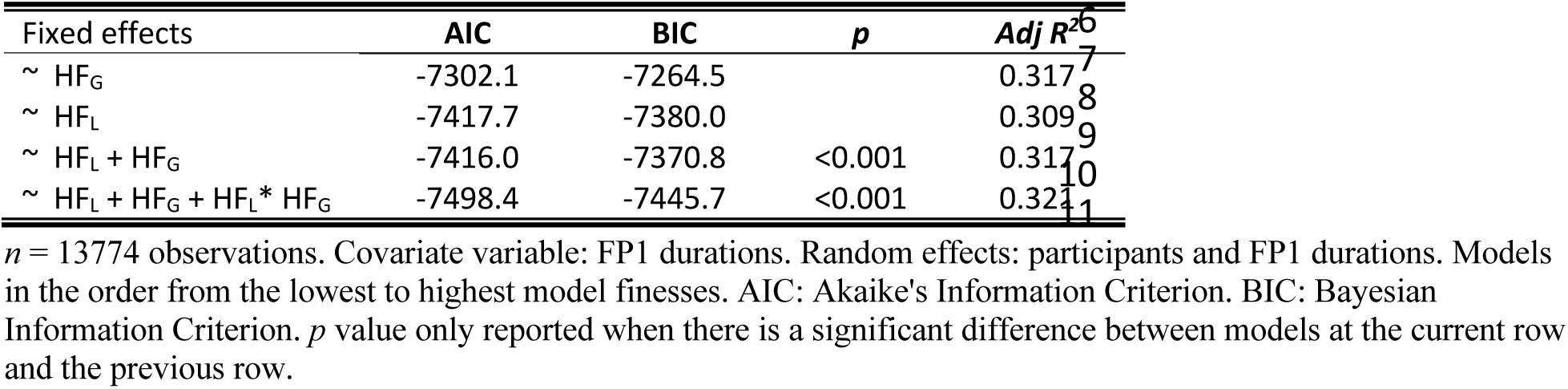
Comparisons of linear mixed-effect models on reaction times following FP2.

**Table 3.** Effects of HF_L_ and HF_G_ on reaction times following FP2.

|  | Estimates | SE | *t value* | *p* | *Adj R*<sup>2</sup> |
| --- | --- | --- | --- | --- | --- |
| (Intercept) | 5.46 | 0.02 | 244.80 | <0.001 | 0.321 |
| HF<sub>L</sub> | -0.21 | 0.02 | -12.98 | <0.001 |  |
| HF<sub>G</sub> | -0.02 | 0.01 | -3.44 | <0.001 |  |
| HF<sub>L</sub> * HF<sub>G</sub> | 0.16 | 0.02 | 9.20 | <0.001 |  |

*n* = 13641 observations.

### Mapping local and global temporal predictions to cortical activations

Building on the behavioral analyses, we further estimated how brain signals encode local and global predictions (HF_L_ and HF_G_) for FP2. Our approach was to identify brain areas where trial-by-trial EEG signals could be reconstructed based on the hazard function over time. To achieve this, we used a forward encoding model, also known as regularized linear regression (Crosse et al., 2016).

In our EEG analysis, to ensure sufficient data length for effective model training, we included trials that lasted longer than 0.7 seconds and did not contain a false alarm. These signals were then segmented, starting from 0.4 seconds after the onset of the warning signal and extending to the onset of the target signal. This segmentation minimized the influence of evoked responses to the warning and target signals. Then, we transformed EEG signals from the channel level (64 EEG channels) to the source level (181*217*181= 7,109,137 brain voxels), using individual 3D electrode locations and structural MR images (see details of the source analysis in Methods). For HF_L_ and HF_G_, values from 0.4 to 2 seconds with a 0.1-second step (10Hz) were splined-interpolated to share the same sampling rate as the EEG source signals (250 Hz) (see Supplementary Figure 1).

To identify which brain areas encode HF_L_, HF_G_, and their interaction, we trained an forward-encoding model to reconstruct the EEG source signals from the hazard values. First, we trained the EEG signals during FP1 on HF_L_ only, which is similar to a typical single foreperiod scenario and was used as a benchmark. Then, we trained the EEG signals during FP2 on (1) HF_L_-only, (2) HF_G_-only, (3) HF_L_+HF_G_, and (4) HF_L_+HF_G_+HF_L_*HF_G_, following the same approach used in the behavioral analysis.

The training was done for each trial and participant with a leave-one-out cross-validation approach. First, one trial was excluded and used as a testing trial, and the remaining trials were used for training (Figure 3A). For EEG signals, the training data comprised dimensions of *n – 1* trials, *m* time points, and 7,109,137 sources (voxels). For the hazard functions, the training data comprised dimensions of *n – 1* trials by *m* time points. Then, we trained the EEG signals from each source on the hazard values with different time lags (76 lags, from -100 to 200 msec with a 4-msec step), resulting in the temporal response function (TRF) (dimensions: 7,109,137 sources and 76 lags) (Figure 3C). Each TRF value represents the weight of the hazard function on the EEG signal at a specific source and lag.

**Figure 3.**
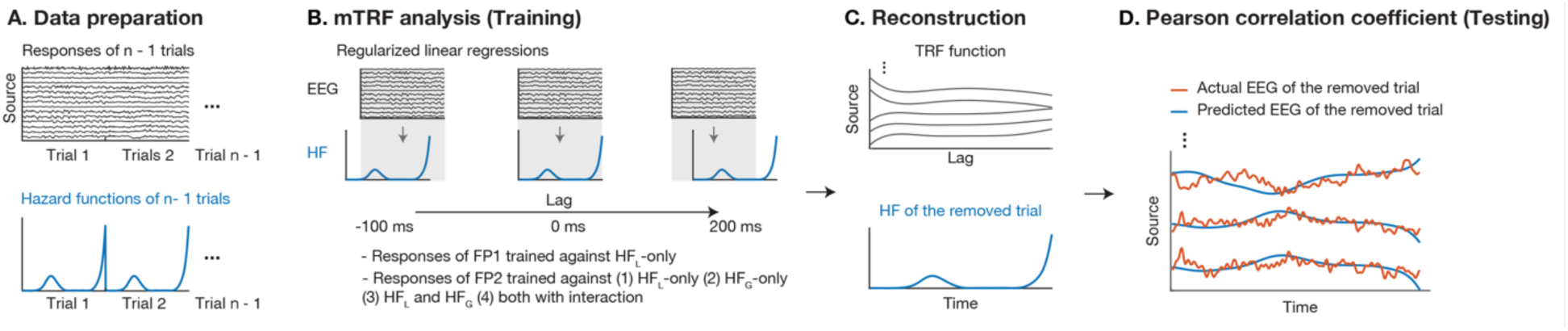
Procedure for the forward-encoding model.

To evaluate the modeling performance, we used the TRF and the hazard functions from the testing trial (Figure 3C, bottom) to reconstruct the EEG signals for each source at a time lag of zero, focusing on the immediate effect of the hazard function on the EEG signals. We then calculated the Pearson correlation coefficients between the reconstructed and actual responses (Figure 3D) and averaged the correlation coefficients across all leave-one-out training (i.e., *n* times from Figure 3A-3D) for each source for each participant. Sources with significant correlations across participants were identified as “significant areas” (n = 31 participants, Monte Carlo method, 1000 permutations, cluster-based correction, two tail, α = 0.05; see more details in Methods).

For FP1, the significant areas included the middle and posterior cingulate areas, superior and middle temporal areas, supramarginal area, calcarine, lingual, and fusiform (Figure 4A). In these significant areas, the TRF values were generally negative (see Supplementary Figure 2), indicating higher HF_L_ values (stronger local predictions) led to smaller responses.

**Figure 4.**
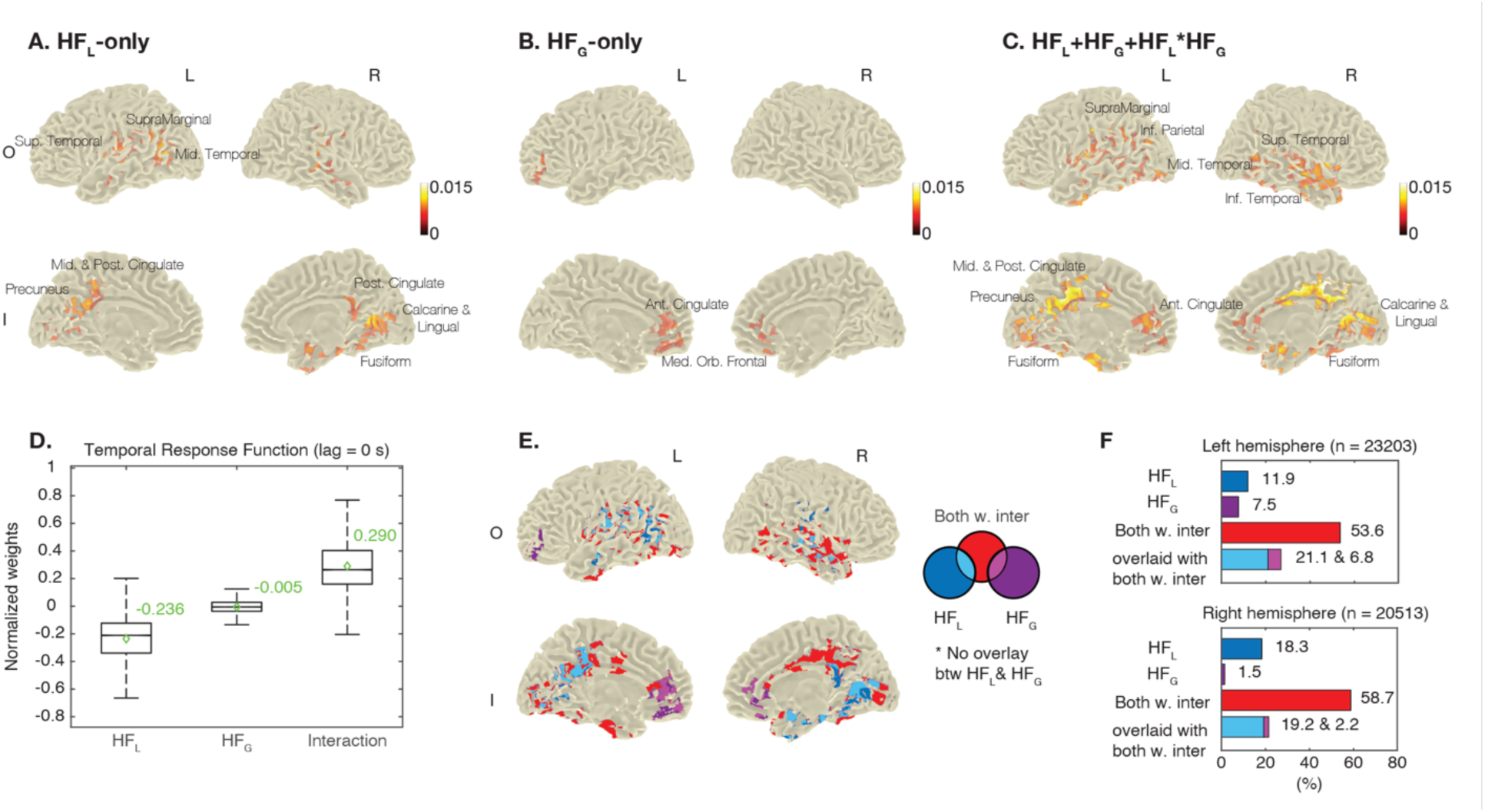
Neural correlates for hierarchical temporal predictions. (A) EEG source responses during FP1 were trained against HF_L_-only. Four cortical surfaces with significant correlation coefficients are shown. The color bar shows correlation coefficients, and the same scale was applied to panels B and C. (B) EEG source responses during FP2 were trained against HF_G_-only, and cortical surfaces with significances are shown. (C) EEG source responses during FP2 were trained against HF_L_+HF_G_+HF_L_*HF_G_, and cortical surfaces with significances are shown. (D) Within the significant areas in the panel C, the box plot shows the positive and negative average TRFs between source responses and hazard values at a 0-second lag. Values were normalized between -1 and 1. (E) The significant areas in the panels A to C were compiled. There is no overlay between the significant areas for HF_L_-only and HF_G_-only. (F) The bar plots show the percentages for the significant areas for each HF over all areas respectively in the left and right hemispheres.

For FP2, significant areas were found only when training with (1) HF_G_-only, and (2) HF_L_+HF_G_+HF_L_*HF_G_. The significant areas for HF_G_-only included the anterior cingulate and medial orbitofrontal areas (Figure 4B). In these areas, we also observed negative TRF values (Supplementary Figure 3), indicating higher HF_G_ values were associated with smaller responses. On the other hand, the significant areas for HF_L_+HF_G_+HF_L_*HF_G_ included the anterior, middle and posterior cingulate areas, superior and middle temporal areas, supramarginal area, inferior parietal area, calcarine, lingual, and fusiform (Figure 4C). This training yielded significantly higher correlation coefficients compared to HF_G_-only (n = 31 participants, Monte Carlo method, 1000 permutations, cluster-based correction, two tail, α = 0.05). This indicates that the EEG signals were better reconstructed when incorporating HF_L_, HF_G_, and their interaction, rather than considering HF_G_ alone, which is consistent with the behavioral model comparisons (Table 2).

For these significant areas for HF_L_+HF_G_+HF_L_*HF_G_, we show the boxplots of the average TRF values for HF_L_, HF_G_, and their interaction (n = 35,104 significant sources for each TRF) (Figure 4D, also see Supplementary Figure 4 for spatial distribution of these values). First, the median TRF values for HF_L_ and HF_G_ were negative, again indicating high local and global predictions are associated with reduced responses. However, the median TRF value for their interaction was positive, suggesting that the influence of HF_L_ on the responses was weaker when the influence of HF_G_ was strong, and vice versa. Again, these findings align with the beta values of the behavioral model.

We further compared the significant areas for HF_L_-only (as shown in Figure 4A), HF_G_-only (Figure 4B), and for HF_L_+HF_G_+HF_L_*HF_G_ (Figure 4C). Here, we refer to these as the HF_L_ area, HF_G_ area, and interaction areas, respectively. In Figure 4E, we found that the HF_L_ area and HF_G_ area (colored separately in dark blue and dark purple) were mutually exclusive. Furthermore, the HF_L_ area and interaction area were broadly overlaid (light blue, the intersection of dark blue and red), while the HF_G_ area and interaction area were shared focally in the prefrontal area (light purple, the intersection of dark purple and red). For the interaction area that were not shared with the HF_L_ and HF_G_ areas (colored in red), several regions were identified, including the middle and posterior cingulate areas, the inferior parietal area, and calcarine.

Figure 4F further illustrates the percentages of these areas in the left and right hemispheres. The interaction area (colored in red) was found to be dominant, accounting for more than half of the total significant areas in both hemispheres. In summary, our findings indicate that the local and global predictions are encoded in distinct brain regions, while the integration of these predictions involves large and additional areas.

## Discussion

Our local-global foreperiod paradigm was designed to create two levels of event timing regularities for temporal prediction. For both behavioral and EEG data, we demonstrated that predictions are made using statistics from both levels and their interactions. We further showed that these processes occur in distinct but overlapping brain regions. Our study establishes an experimental and analytical platform to isolate latent dynamics in hierarchical temporal prediction, a crucial step towards unifying predictive coding of “what” and “when” information.

### Prefrontal cortex involvement in processing long-scale information

Our investigation used a forward encoding model to estimate how the brain encodes hierarchical temporal predictions based on learning the local and global regularities. The hazard function is a representation of a prediction updated over time, and this dynamic characteristic was kept and assigned as a time-resolved regressor in model training. By training against HF_L_ and HF_G_ values, we disentangled their corresponding neural correlates and extracted the correlates of their interaction at the source-level areas.

Specifically, only when the sequence structure was considered (the global regularity), the neural correlates in the anterior cingulate and medial frontal areas were identified. This prefrontal involvement has been reported in previous neuroimaging studies showing differences when comparing responses in the variable foreperiod paradigm (i.e., a uniform distribution of the foreperiod) and in the fixed foreperiod paradigm (Coull et al., 2016; Vallesi, McIntosh, Shallice, et al., 2009; Vallesi, McIntosh, & Stuss, 2009). Additionally, in rodents, inactivations of bilateral dorsomedial prefrontal cortex (dmPFC) led to premature motor responses, indicating top-down inhibition on the motor cortex during preparation (Narayanan et al., 2006; Narayanan & Laubach, 2006). While these previous studies localized prefrontal activity by manipulating single foreperiod occurrences (akin to the local regularity in our design), we observed similar activity only in processes of the foreperiod sequence (the global regularity). In fact, increased responses were found in ERPs during both FP1 and FP2, and a significantly positive waveform at the frontal electrodes was found by comparing ERPs during FP1 and FP2 (Supplementary Figure 5). We suggest that the frontal area provides sustained monitoring. For instance, in the previous studies, increased prefrontal activity was found in the condition of unimodal probability distributions, compared to precisely invariant FP. Also, the retention of temporal information over a longer scale is supported by prefrontal neurons activating differentially to store different durations of the first and second cues (Oshio et al., 2006). For our findings, ramping-up prefrontal activity may reflect tracking of FP1 and FP2 in the two-foreperiod sequence.

### A distinct brain network for the integration of hierarchical temporal predictions

We used the multiplication of the HF_L_ and HF_G_ values to represent the integration of local and global predictions. This approach was inspired by the interaction term in regression models, which allows for the combined effect of two or more dependent variables on the independent variable, in addition to their individual main effects. Interestingly, in both behavioral regression and neural encoding models, we observed a positive effect of the interaction term in both behavioral and neural models. This indicates that the prediction at one level influences the reaction time and brain response more strongly when the prediction at the other level is weak, and vice versa. Furthermore, a wide range of brain areas were found to process integration, as there were no shared areas between significant correlates to HF_L_ and HF_G_. Particularly, the posterior cingulate cortex was notably identified with a large distribution among the important areas involved in the interaction. The posterior cingulate cortex was part of the frontoparietal network comprising precuneus and inferior frontal gyrus, and functional connectivity of this network mainly increased with update values in an ideal Bayesian observer model. The update is quantified as a divergence between prior and posterior probabilities of the current FP, compared to surprise, inversely and non-linearly correlated with the hazard function (Visalli et al., 2019, 2021). Furthermore, the neural correlate of integration localized preferentially at the right side of posterior cingulate cortex further strengthens a link to the top-down updates. Hemispheric lateralization in temporal processes has been proposed, where the right hemisphere would be better at learning previous information to predict future onset while the left hemisphere would be better at comparisons between a test stimulus interval and a target stimulus interval (Coull & Nobre, 2008). We infer that simultaneously processing multiple levels of prior information would require intense prediction updates in the posterior cingulate cortex.

On the other hand, the inferior parietal area was identified though its distribution was relatively sparse in our results. It has been shown that neural activity in the lateral intraparietal neuron field changed differently in blocks of unimodal and bimodal distributions (Janssen & Shadlen, 2005). Analogously, we created different bimodal distributions of the local regularity (e.g., 50-50) and the global regularity (e.g., 80-20), and discrimination between the probability distributions may also be a function of the integration hub. Here, it is worth noting that Janssen and Shadlen adopted a smooth version of HF (called temporal-blurred HF), assuming the precision of timing decreases as time goes on. Also, recent studies using a probabilistically blurred PDF instead of a HF provided a better explanation for the reaction time (Grabenhorst et al., 2019, 2021). To thoroughly examine possible representations of the temporal predictions, we applied different smooth versions of HF and PDF to the behavioral and neural models but did not observe significant improvements (Supplementary Method and Supplementary Tables 3 and 4). This lack of improvement may result from a lower quality of trial-by-trial EEG signals compared to neuronal recording, making it challenging to capture differences between the smooth versions (Herbst et al., 2018).

While our study provided insights into brain networks involved in hierarchical temporal predictions based on time-series responses, it is also critical to understand how these networks function and communicate within the oscillatory domain. The alpha/beta oscillation has been suggested as a representation of the top-down prediction signal, while the gamma oscillation is associated with the bottom-up prediction error signal (Bastos et al., 2020). This aligns with findings that show that alpha power prior to a target changed with the hazard function (Herbst & Obleser, 2017). Furthermore, local and global predictions of “what will happen” in a sequential context have been shown in different ranges of the beta oscillation (Chao et al., 2022). To further investigate this frequency ordering of hierarchical predictions of “when an event will happen”, we should combine time-frequency analysis as well as functional connectivity analysis in our future work.

### The asymmetrical sequential effect observed in bimodal distributions

While our results demonstrated that reaction times were faster as hazard values were higher, we also observed an asymmetrical sequential effect due to our design composed of long/ short FP1 paired with long/ short FP2. The effect describes that reaction times increase if the preceding foreperiod trial is longer than the current one in the variable foreperiod paradigm. A model based on the principle of trace conditioning has been proposed to account for the effect (Los & van den Heuvel, 2001). Initially, in the case of a uniform probability distribution, the chance of a target to occur at any time is equal and thus the associated weights for each critical moment are equal. During the preparation for target occurrence, weights decrease as the corresponding critical moment passes (called extinction). This so-called extinction becomes weaker as time elapses, meaning that the weight associated with a late moment decreases less or remains relatively unchanged. At the imperative moment when the target is presented, the response is made, leading to an increase in the weight associated with the imperative moment (reinforcement). Then, the associate weights are passed to the next foreperiod trial. If the imperative moment is shorter (i.e., a target occurs after a shorter foreperiod than the previous one), the associated weight was already weaker for that moment, thus resulting in a slower response.

Regarding the asymmetrical sequential effect, two points should be noted: (1) Our regression results still show modulation from the hazard function while durations of FP1 were controlled as a covariate, and (2) Our forward encoding model results excluded the sequential effect. Regarding (1), we calculated the average reaction times following long and short FP2 after either short or long FP1 (i.e., *LL*, *LS*, *SS*, and *SL*) for four blocks (Supplementary Figure 6, the visualization of Supplementary Table 2). The asymmetry effect was observed, where reaction times following FP2 tended to be longer when FP1 had a longer duration (*L<u>S</u>* vs. *S<u>S</u>*), but the effect were different among the blocks due to our manipulations for different probability distributions. For example, in Supplementary Figure 6, the asymmetry between the reaction times following FP2 in *L<u>S</u>* and *S<u>S</u>* is smaller in Block 4 but larger in Block 3. This is because Block 4 composed of more trials of *LS* (40%) than Block 3 (10%), and fewer trials of *SS* (10%) than Block 3 (40%). Therefore, to carefully identify the effect of the foreperiod on reaction time (i.e., temporal hazards) and exclude the sequential effect, we controlled the FP1 length in our regression analysis.

Regarding (2), it is known that anatomical locations responsible for the effect of the foreperiod on reaction and the sequential effect are distinct (Vallesi et al., 2007). Patients with lesions to the right frontal area had a weaker FP-RT effect while the sequential effect remained intact. In contrast, lesions to the left premotor area diminished the sequential effect. Here, we adopted the forward encoding model and identified the neural correlates of HFs mostly in the posterior cingulate and frontal areas; however, an involvement of the premotor area was not detected. This suggests the forward encoding model as a plausible analysis tool to extract responses of temporal predictions while eliminating confounds due to the sequential effect. Still, although we used individual MRI images to reduce the volume conduction effect and reconstruct source-level signals, we should carefully consider the spatial limitations of the EEG when coming to a conclusion. When using functional magnetic resonance imaging (fMRI), it is also important to consider the trade-off cost between losing dynamics characteristics of temporal predictions changing over time and gaining more precise spatial information.

In conclusion, to our knowledge, this study is the first to reveal how the brain integrates multi-level information of event timing for prediction establishment. This also supports the hierarchical organization of the predictive-coding theory in the “when” domain. Since we solely manipulated the event timing, in order to generalize this well-known and fundamental theory, we plan to simultaneously manipulate multi-level information in both the “what” domain (e.g., tone pitches) and the “when” domain (e.g., tone onset) in future research.

## Method

### Participants

We recruited 34 participants (17 males and 17 females; age: 23 ± 2.9 years old). The inclusion criteria were: (1) age between 20 and 50 years old; (2) no severe deficit in hearing loss, eyesight, and color discrimination that could cause problems in understanding experimental procedures; (3) no medical history and diagnosis of neurological or psychological diseases reported by the participant. All participants signed consents after understanding the procedure and before the experiment started. The protocol was approved by the ethical committees of the University of Tokyo (No. 21-372). The data of three participants were excluded because they were easily distracted during the experiment or missed structural MRI acquisition (16 males and 15 females; age: 23 ± 3 years old).

### Stimulus

The warning signal was composed of an auditory stimulus (three combined pitches: 350, 700, and 1400 Hz, 55 dB) and a visual stimulus (a white dot with a 15-pixel diameter placed at the center). The duration of the warning signal was 0.1 second. The target signal was composed of an auditory stimulus (three combined pitches: 500, 1000, and 1500 Hz) and a visual stimulus (a white dot with a 15-pixel diameter placed at the center). The duration extended until the *zero* button was pressed or for 1 second if no *zero*-pressed response was detected. The stimulation layout on the monitor (a resolution of 1920*1080 pixels) included (1) a gray background (RGB: [85,85,85]), (2) a fixation (a white cross with a 40-pixel diameter) placed 110 pixels below the center of the monitor. The auditory stimuli were delivered through a pair of desktop speakers. The stimulation was programmed using MATLAB-based Psychtoolbox (Kleiner et al., 2007; Pelli, 1997) and presented in a dim and sound-proof booth.

### Local-global foreperiod sequence paradigm

We paired two foreperiod trials as a sequence (FP1 and FP2) to establish local and global regularities. With each sequence, two foreperiod trials were separated by a 1.2-second interval between the offset of the target in FP1 and the onset of the warning in FP2. Between consecutive sequences, the interval ranged from 3 to 3.2 seconds, with increments of 0.05 seconds, between the offset of the target in FP2 and the onset of the warning in the next FP1. To make foreperiod chunking obvious to the participant, a black square (a size of 400*400 pixels placed 110 pixels below the center) was presented 0.5 seconds after the press response for the target in FP2 or 1 second after the onset of the target when the press response was absent. Foreperiod trials within the duration range between 0.4 and 1.1 seconds and between 1.3 to 2 seconds are denoted as *S* and *L*, respectively. The participants underwent prior testing to be familiar with foreperiod trials having relatively short or long durations. There were four sequence types: *LL*, *SS*, *LS*, and *SL*, where the first and the second in each pair represent the duration of FP1 and FP2 respectively.

Four blocks were designed, each containing 50 trials of two-foreperiod sequences. In Block 1, there were 25 trials of *LL*, and 25 trials of *SS*, leading to PDF_L_ with two non-overlay unimodal probability distributions respectively peaking at 0.8 and 1.6 seconds. The PDF_L_ with a 50-50 ratio (referred to as the 50-50 PDF_L_) represents 50% of cumulative probabilities in the range of *S* and 50% of cumulative probabilities in the range of *L*. For PDF_G_, a 0-100 distribution after long FP1 was calculated, representing close to 0% of cumulative probabilities in the range of *S* and close to 100% of cumulative probabilities in the range of *L*. Similarly, a 100-0 distribution after short FP1 was calculated. In Block 2, there were 25 trials of *LS* and 25 trials of *SL*, resulting in a 50-50 PDF_L_, a 100-0 PDF_G_ after long FP1, and a 0-100 PDF_G_ after short FP1. In Block 3, there were 20 trials of *LL*, 5 trials of *LS*, 20 trials of *SS*, and 5 trials of *SL*, resulting in a 50-50 PDF_L_, a 20-80 PDF_G_ after long FP1, and an 80-20 PDF_G_ after short FP1. In Block 4, there were 5 trials of *LL*, 20 trials of *LS*, 5 trials of *SS*, and 20 trials of *SL*, resulting in a 50-50 PDF_L_, an 80-20 PDF_G_ after long FP1, and a 20-80 PDF_G_ after short FP1. Supplementary Figure 7 detailed trial numbers of foreperiod with different durations in the four blocks

During the experiment, the participants were instructed to (1) sit comfortably, (2) rest their chins on a chin supporter, (3) look at the fixation to minimize head and eye movements, and (4) press the *zero* button as soon as the target signal appeared. The four blocks were presented in a random order and two subsequent repeats followed. After each block presentation, there was a short rest, and the participants were asked to identify which sequence type appeared frequently.

### Data acquisitions

We recorded EEG signals using the 64-channel actiCAP slim from Brain Products and captured individual electrode locations using a 3D camera (brand: STRUCTURE). To reconstruct EEG sources, we also collected individual structural MRI images (T1) using SIEMENS 3T Magnetom Prisma. These two measurements were conducted either on the same day or on two separate days.

#### EEG recording

The 64-channel electrode cap was capped according to anatomical landmarks including nasion, left and right porions (a top of the ear canal). We captured electrode locations and the three landmarks (labeled with red stickers) using the 3D camera mounted on an iPad. During EEG recording, event codes were sent at the onset of the warning signal in each foreperiod trial. Raw EEG signals were recorded with a 1000-Hz sampling rate.

#### MRI acquisition

Before image acquisition, the participants completed an MRI safety checklist, and a device was used to detect whether there was any metal material on body. The three anatomical landmarks were labeled using sugar candies (小林製薬のブレスケア) which can be identified in MRI images. The use of the three landmarks allowed us to align electrode locations and MRI images to the same anatomical axis in later analyses. During acquisition, comfortable air-filled paddings were used to prevent severe head motion, and earplugs were used to reduce scanner noise. The T1 acquisition protocol included a field of view: 240 × 256 mm, 300 × 320 matrix, TR: 2400 ms, TE: 2.22 ms, flip angle: 8◦, and 0.8-mm slice thickness.

## Data analyses

### Hazard function

HF, describing the conditional probability of an event occurring at time *t* given it has not yet occurred, was used to estimate dynamics of temporal predictions. The formula is listed below:

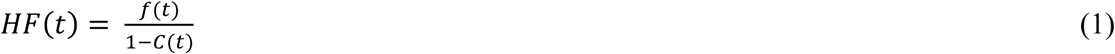

where *t* represents time points with the range of 0.4 to 2 seconds. The function *f* represents the probability distribution of the foreperiod in a block. The function *C* represents the cumulative probabilities until time *t*. When 1 − *C*(*t*) is close to zero, *HF*(*t*) is toward infinity. We replaced the infinite value with the maximum hazard value before time *t*. Finally, we normalized hazard values between 0 to1.

### Linear mixed-effect analysis

To estimate correlations between hazard values (i.e., temporal prediction) and reaction times, we used linear mixed-effect models while individual differences were controlled. The durations of FP1 (*L* or *S*) were also controlled when reaction times following FP2 were modeled. The regression models are as below:

For FP1

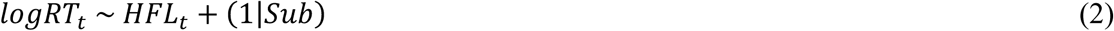

For FP2

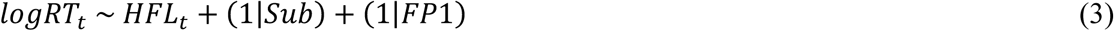

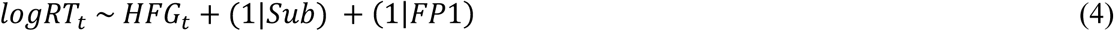

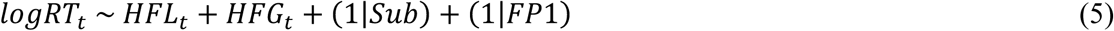

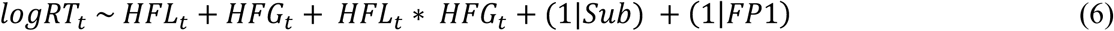

where *logRT_t_* represents log-transformed reaction times to the target signal appearing at time *t* (relative to the onset of a warning signal); *HFL_t_* represents a HF_L_ value at time *t*; *HFG_t_* represents a HF_G_ value at time *t*.

The analyses were conducted using R (4.2.2). The functions, *lmer*, *anova*, and *r.squaredGLMM*, were used for the regression analyses, model comparisons, and adjusted R-squared, respectively.

### EEG preprocessing

The raw EEG signals were preprocessed using MATLAB-based EEGLAB for each participant. Firstly, channels showing high amplitudes (over 100 microvoltage) in over than half of the dataset were removed (*pop_select.m*), and signals below 0.1 Hz were filtered out (*pop_eegfiltnew.m*). Independent component analysis (ICA) was used for component extraction (*pop_runica.m*), and components containing artifacts were removed using the ADJUST plugin toolbox (*interface_ADJ.m*). Next, the data was segmented into epochs with a time range between -0.8 and 2.8 seconds relative to the onset of the warning for each foreperiod trial (*pop_epoch.m*), and epochs showing high amplitudes (over 100 microvoltage) were manually removed. The epoch data was then referenced to the average of signals across the 64 channels (*pop_reref.m*), and 1-Hz low-pass and 55-Hz high-pass filters were applied. Finally, the data was baseline-corrected (-0.2 and 0 seconds; *pop_rmbased.m*) and down-sampled to 250 Hz (*pop_resample.m*).

For further analyses, we disregarded trials shorter than 0.8 seconds or trials with the false alarm. The processed EEG signals were estimated between 0.4 seconds after the onset of the warning and the onset of the target.

### Source reconstruction analysis

The analysis aims to estimate volume conductivity (i.e., forward model) and interpolate sources of electrical potentials (i.e., inverse model) for each participant. First, a head model (also known as a volume conduction model) was created using structural MRI images with the finite element method. The images were resliced into isotropic dimensions (256 * 256 * 256 [voxel]) (*ft_volumeslice.m*) and realigned to the three candy-labeled markers (*ft_realign.m*). Then, the images were segmented into five tissue types (gray matter, white matter, cerebral cerebrospinal fluid, skull, and scalp) (*ft_volumesegent.m*). The boundary was adjusted to create a hexahedron for each voxel with the shifting parameter set to 0.3 (*ft_preparemesh.m*). The volume conductivity, [0.33 0.14 1.79 0.01 0.43], was assigned in the order of the above-mentioned tissue types (*ft_prepare_headmodel.m*). Second, the electrode locations captured in the 3D image were aligned according to the three red-dot-labeled markers (*ft_meshrealign.m*), and electrode names were assigned manually (*ft_electrodeplacement.m*). The locations were further adjusted according to the brain shape for better fitness (*ft_plot_headshape.m*; *ft_electroderealign.m*).

Third, the *leadfield* (potential contributions from dipoles to electrodes) was created with the head model and the electrode locations, with a resolution of 1 center (*ft_prepare_leadfield.m*). Then, linear-constraint minimum-variance beamforming (*lcmv* beamforming) was applied to extract spatial filters (*ft_sourceanalysis.m*). The input products included the processed EEG signals, the head model, and the *leadfield*. The output product was spatial filters (dimensions: *xyz-axes* * *electrode * source*). To acquire source signals in dimensions of *time points* and *source*, spatial filters of each source (*xyz-axes* * *electrode*) were multiplied by the EEG (*electrode* * *time points*), resulting in a time course in three axes (*xyz-axes* * *time points*). We used principal component analysis to extract a time course along a dominant axis (*time points*).

### Multivariate Temporal Response Function analysis (mTRF)

The multivariate Temporal Response Function analysis (mTRF) was used to evaluate correlations between hazard values (i.e., temporal prediction) and EEG source signals. This analysis enables a convoluted regression of a stimulus vector (e.g., HF_L_) against neural responsess (e.g., a time course of a source), and has been widely used in speech research (Di Liberto et al., 2015) and recently in time research (Herbst et al., 2018). The formula is as follows.

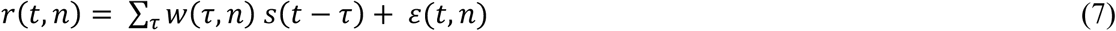

Where *r* represents the EEG source response for one participant. *t* represents a time point. *n* represents an EEG source. τ represents a time lag. *w* represents the temporal response function. *s* represents the hazard function. ω represents a residual response not explained by the model. The analysis was conducted using the MATLAB-based mTRF toolbox. Parameters were set to -0.1 and 0.2 for minimal and maximal time lags, and 1 for *lamba*. We determined these parameters based on optimal correlation coefficients between hazard values and EEG signals (*mTRFcrossval.mat*). To match the sampling rate of the source response (in 250 Hz), hazard values (in 10Hz) were splined-interpolated (*interp1.mat*).

For each participant, the leave-one-out correlation coefficients were evaluated as follows (Figure 3): (1) for each trial, it was temporarily removed from the data; (2) for each source, responses of the remaining (n−1) trials were trained against the corresponding hazard values (*mTRFtrain.m*) (3) The output was the temporal response function *w*; (4) for each source, the temporal response function and the hazard function of the removed trial were used to reconstruct a predicted response of the removed trial; (5) Pearson correlation coefficient between the predicted response and the actual response was estimated for each source (*mTRFpredict.m)*. For FP1, the response was trained against HF_L_. For FP2, the response was trained against (1) HF_L_-only, (2) HF_G_-only, (3) HF_L_ and HF_G_, and (4) both with the interaction term. For each participant, the final correlation coefficients (*n* trials * 7109137 sources) were averaged across trials.

Moreover, we also trained the response against shuffled hazard values as a benchmark control. We shuffled hazard values for each trial and participant. We segmented hazard values into five parts (e.g., 250 values with 50 values per part) and rearranged the order of these parts. This shuffle can contain partial changes of the hazard function, compared to the total randomization. For example, one shuffle with the order of 2-1-3-5-4 would thus be (51th-100th)-(1st-50th)-(101st-150th)-(201st-250th)-(151st-200th). For the shuffled HF as controls, we only found significant correlates of shuffled HF_L_ (Supplementary Figure 8) and disregarded them from the correlates shown in Figure 4A.

### Statistics and visualization

In order to assess whether the correlation coefficients significantly differ from zero, we performed group-level analyses, and the procedure included source interpolation, volume normalization, and source statistics. For each participant, sources, each containing the correlation coefficient, were interpolated back to their individual MRI images (*ft_sourceinterpolate.m*). The anatomical layout was then normalized to the standard brain map, with parameters set to T1.nii in SPM12 and a non-linear transformation (*ft_volumenormalise.m*). Significances across the participants were tested, with parameters set to Monte Carlo method, cluster-based correction, 1000 randomization, a two-tailed test, and an alpha level of 0.05 (*ft_sourcestatistics.m)*. The significances were visualized on cortical surfaces of the MNI brain with parameters set to a nearest projection and no lighting (*ft_sourceplot.m*).

## Supporting information

All supplementary materials

## Data availability

The raw data and scripts will be shared in public after publication.

## Contributions

Z.C.C. and Y.T.H conceptualized the study. Y.T.H collected the data and performed data analysis. Y.T.H wrote the paper, and Z.C.C edited the paper. The two authors contributed to and approved the final paper.

## Acknowledgments

We thank Lu Li and Junko Taniai for helping with participant recruitment and experiment preparation. We also thank Felix B. Kern for proofreading. This study was supported by Japan Society for the Promotion of Science, Japan (to Y.T.H) and World Premier International Research Center Initiative (WPI), MEXT, Japan (to Z.C.C.).

