## Supplementary material for "Temporal Prediction through Integration of Probability Distributions of Event Timings at Multiple Levels": All supplementary materials

**Supplementary Method**

There were other representations of predicting when an event to occur, in addition to the hazard function (HF), including original probability distribution, temporally blurred PDF, and probabilistically blurred PDF, and their transformation to HF (Grabenhorst et al., 2019, 2021). First, although HF can exactly represent dynamic updates of temporal predictions, its continuous and complicated computations over time have been challenged, and PDF as the foundation of HF has been proposed for temporal prediction representations. Second, uncertainty in estimating probabilities increases over time. In other words, the brain predicts a stimulus onset less precisely as time elapses. This phenomenon is termed the temporally blurred effect, where uncertainty is modeled by a Gaussian function with the standard deviation scaling with time. The equations for temporally blurred PDF and temporally blurred HF are listed below.

$$f_{tb}(t) = \frac{1}{\varphi t\sqrt{2\pi}} \int_{-\infty}^{\infty} f(\tau) \cdot e^{-(\tau - t)^2/\,(2\varphi^2 t^2)} d\tau \qquad (1)$$

$$HF_{tb}(t) = \frac{f_{tb}(t)}{1 - C_{tb}(t)} \qquad (2)$$

where *t* represents time points, ranging from 0.4 to 2 sec. The function $f_{tb}$ represents the temporal-blurred probability distribution of the FP in a block. $t \cdot \varphi$ represents the standard deviation scaling with time. In the current study, we tested $\varphi$ from 0.15 to 0.35 and set it at 0.22 based on its better fitness for RTs. The parameter was also close to previous studies (Grabenhorst et al., 2019, 2021; Janssen & Shadlen, 2005). The function $C_{tb}$ represents the accumulation of temporal-blurred probabilities up to time *t*. When $1 - C_{tb}(t)$ approaches zero, $HF_{tb}(t)$ becomes an infinite value, and we replaced the infinite value with the maximum hazard value before time *t*. Subsequently, hazard values were normalized between 0 to1.

Third, uncertainty can be also scaled with precision, meaning that the brain predicts a stimulus onset less precisely due to the lower chance (probability) at that specific time. This is termed the probabilistically blurred effect, where uncertainty is modeled by a Gaussian function with the standard deviation scaling with probability.

$$\sigma_{min} = \varphi \cdot t_{min} = \varphi \cdot 0.4 \tag{3}$$

$$\sigma_{max} = \varphi \cdot t_{max} = \varphi \cdot 2.0 \tag{4}$$

$$s(t) = \left[1 - \frac{(f(t) - f_{min})}{(f_{max} - f_{min})}\right] \cdot (\sigma_{max} - \sigma_{min}) + \sigma_{min} \tag{5}$$

$$f_{pb}(t) = \frac{1}{s(t)\sqrt{2\pi}} \int_{-\infty}^{\infty} f(\tau) \cdot e^{-(\tau - t)^2 / (2s(t)^2)} d\tau \tag{6}$$

where $\varphi$ was set to 0.22 as well, based on its better fitness for RT. The function $s$ represents the standard deviation scaling with probability, where a higher probability leads to a lower standard deviation and vice versa. The function $f_{pb}$ represents the probabilistically blurred probability distribution of the FP in a block. The transformation from $f_{pb}$ to the probabilistically blurred HF is the same as the formula (2).

### Supplementary Table 1. Average reaction times

| logRT | Block 1 | Block 2 | Block 3 | Block 4 |
|---|---|---|---|---|
| FP1 | 5.419 ± 0.103 | 5.421 ± 0.112 | 5.431 ± 0.120 | 5.425 ± 0.120 |
| FP2 | 5.384 ± 0.126 | 5.438 ± 0.123 | 5.405 ± 0.131 | 5.436 ± 0.123 |

Mean ± Standard deviation.

### Supplementary Table 2. Average reaction times following FP2

| logRT | *After long FP1* | | *After short FP1* | |
|---|---|---|---|---|
| | *L (LL)* | *S (LS)* | *S (SS)* | *L(SL)* |
| Block 1 | 5. 370± 0.132 | | 5. 396± 0.126 | |
| Block 2 | | 5. 473± 0.125 | | 5. 399± 0.137 |
| Block 3 | 5. 375± 0.137 | 5. 510± 0.136 | 5. 411± 0.142 | 5. 394± 0.115 |
| Block 4 | 5. 397± 0.132 | 5. 481± 0.130 | 5. 450± 0.114 | 5. 391± 0.144 |

Mean ± Standard deviation. *LL*: RT to long FP2 after long FP1. *LS*: RT to short FP2 after long FP1. *SS*: RT to short FP2 after short FP1. *SL*: RT to long FP2 after short FP1.

### Supplementary Table 3. Comparisons of linear mixed-effect models on reaction times following FP1

| Fixed effects | AIC | BIC | *p* | *Adj $R^2$* |
|---|---|---|---|---|
| **~ $HF_L$** | **-8148.5** | **-8118.4** | | **0.292** |
| ~ $Prob_L$ | -7914.6 | -7884.5 | | 0.277 |
| ~ Temp-blurred $HF_L$ | -8158.7 | -8128.6 | | 0.292 |
| ~ Temp-blurred $Prob_L$ | -8047.7 | -8017.7 | | 0.287 |
| ~ Prob-blurred $HF_L$ | -8196.0 | -8165.9 | | 0.295 |
| ~ Prob-blurred $Prob_L$ | -7963.9 | -7933.8 | | 0.280 |

*n* = 13641 observations. Random effect: participants. AIC: Akaike's Information Criterion. BIC: Bayesian Information Criterion. *p* value only reported when there is a significant difference between models at the current row and the previous row. Adj $R^2$: adjusted R-squared. Prob: probability distribution. Temp-blurred: temporally blurred. Prob-blurred: Probabilistically blurred.

### Supplementary Table 4. Comparisons of linear mixed-effect models on reaction times following FP2

| Fixed effects | AIC | BIC | *p* | *Adj $R^2$* |
|---|---|---|---|---|
| **~ $HF_L$ : $HF_L$** | **-7498.4** | **-7445.7** | | **0.321** |
| ~ $Prob_L$ : $Prob_G$ | -7317.0 | -7264.3 | | 0.310 |
| ~ Temp-blurred $HF_L$ : Temp-blurred $HF_G$ | -7632.0 | -7579.3 | | 0.329 |
| ~ Temp-blurred $Prob_L$ : Temp-blurred $Prob_G$ | -7453.8 | -7401.1 | | 0.319 |
| ~ Prob-blurred $HF_L$ : Prob-blurred $HF_G$ | -7621.0 | -7568.3 | | 0.329 |
| ~ Prob-blurred $Prob_L$ : Prob-blurred $Prob_G$ | -7347.6 | -7294.9 | | 0.311 |

*n* = 13774 observations. Covariate variable: FP1 durations. Random effects: participants.

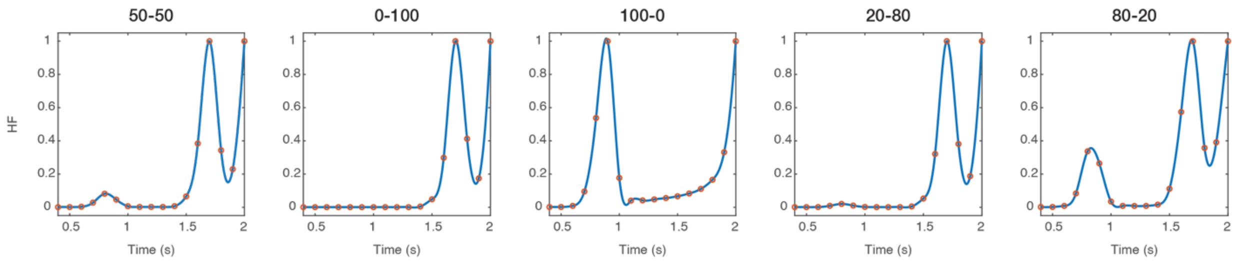


**Supplementary Figure 1 Splined-interpolated hazard values.** The original hazard values at a 10-Hz sampling rate are represented in the orange while the interpolated values at a 250-Hz sampling rate are represented in the blue.

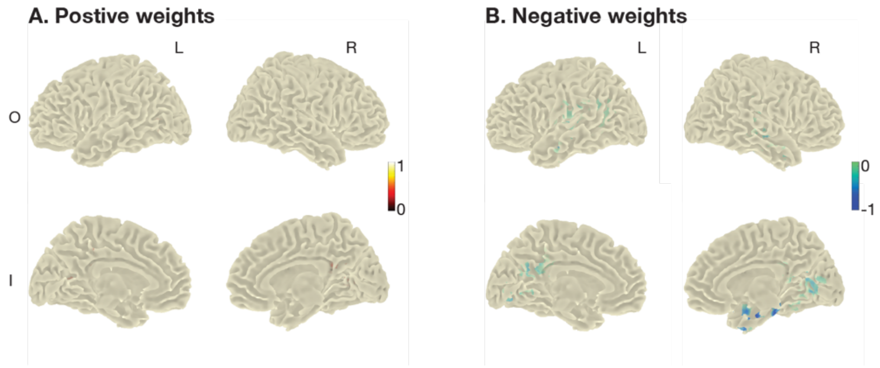


**Supplementary Figure 2 Temporal response function of $HF_L$.** Within the significant area for $HF_L$-only in Figure 4A, the positive and negative average TRF (i.e., weight) between source responses and $HF_L$ values at a 0-second lag are shown. Values were normalized between -1 and 1.

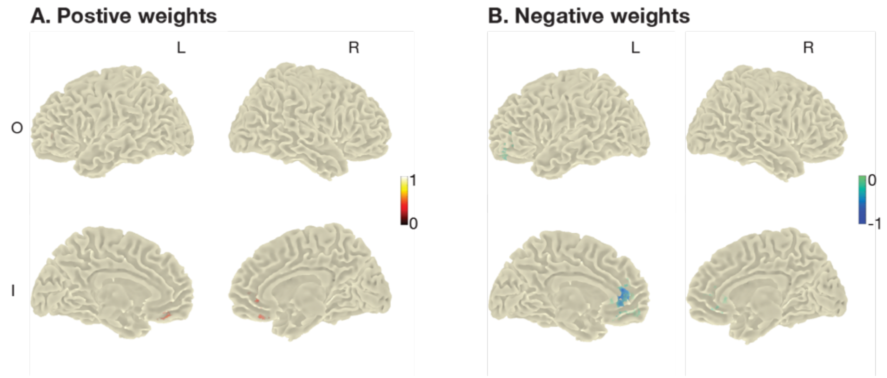


**Supplementary Figure 3 Temporal response function of $HF_G$.** Within the significant area for $HF_G$-only in Figure 4B, the positive and negative average TRF between source responses and $HF_G$ values at a 0-second lag are shown. Values were normalized between -1 and 1.

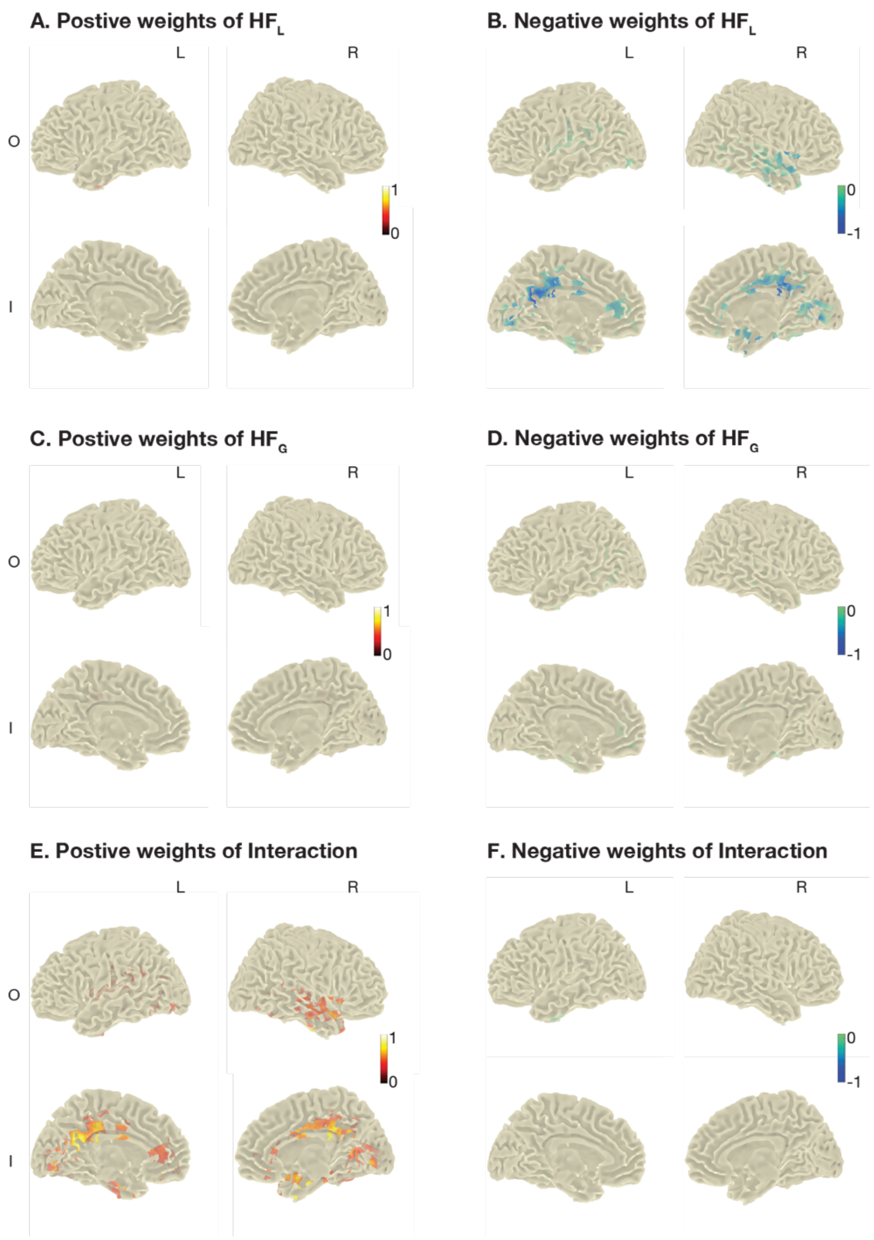


**Supplementary Figure 4 Temporal response functions of $HF_L$, $HF_G$, and their interaction.** Within the significant area for $HF_L+HF_G+HF_L*HF_G$ in Figure 4C, the positive and negative average TRFs between source responses and each of those three values at a 0-second lag are shown. Values were normalized between -1 and 1.

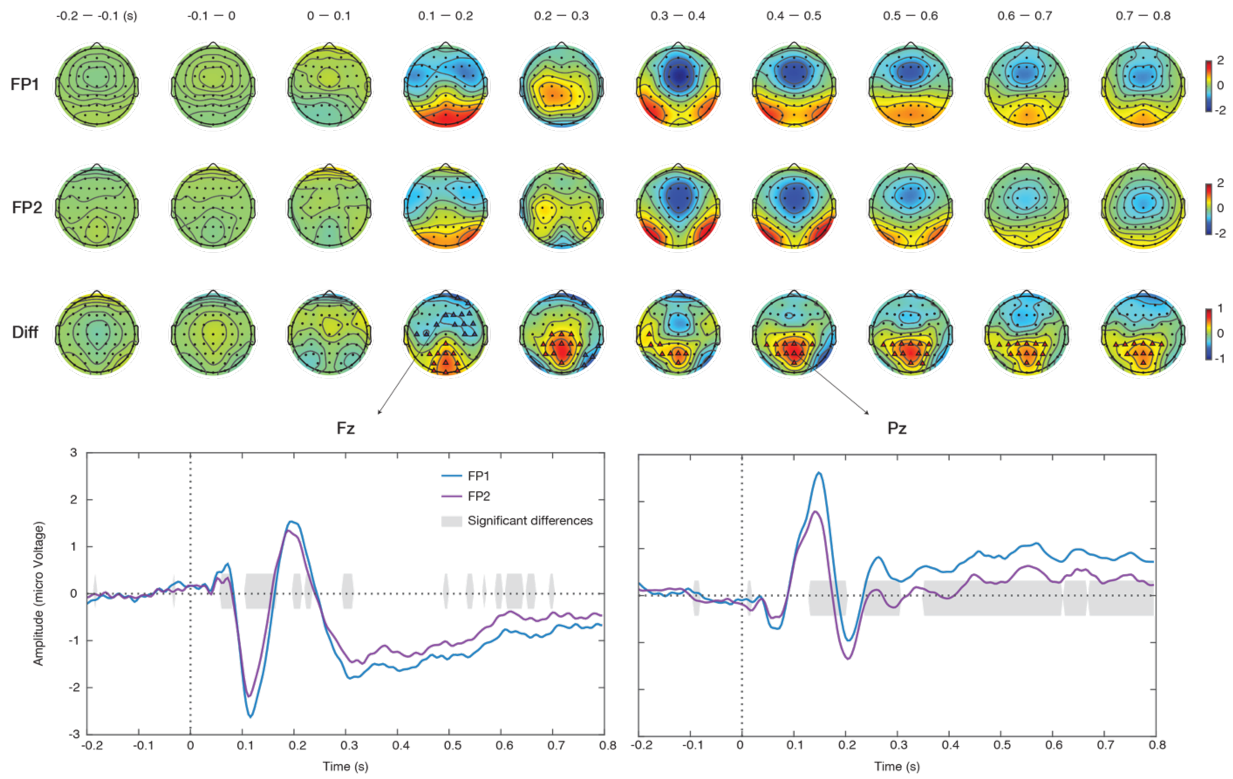


**Supplementary Figure 5 Event-related potentials (ERP) during FP1 and FP2.** We averaged processed EEG signals in FP trials lasting longer than 0.8 second. Time zero represents the onset of the warning signal. Differences between ERPs during FP1 and FP2 (FP1 — FP2) were tests using the Monte Carlo method and cluster-based correction (1000 randomization, a two-tailed test, and an alpha level of 0.05). Triangles in the topography and gray shades in the time course represent the significances

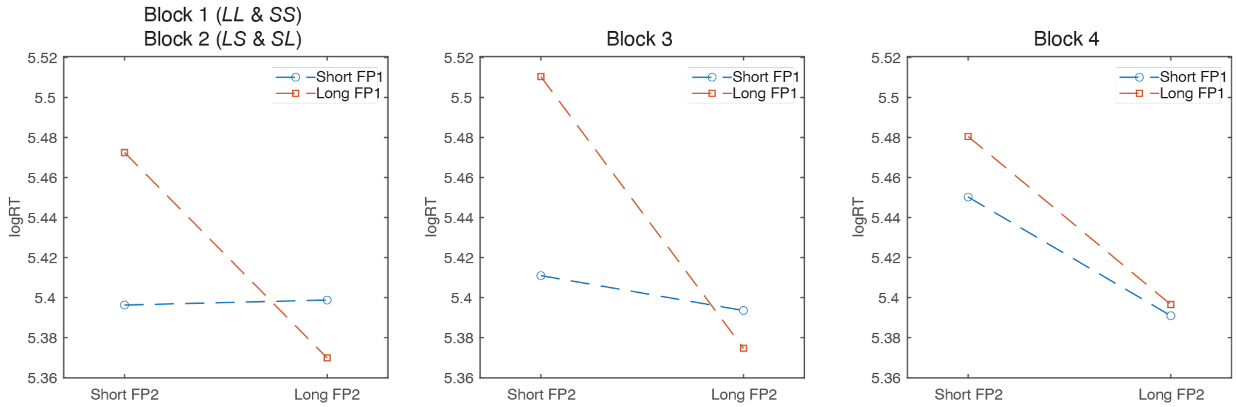


**Supplementary Figure 6 The average reaction time following FP2.** The visualization of Supplementary Table 2

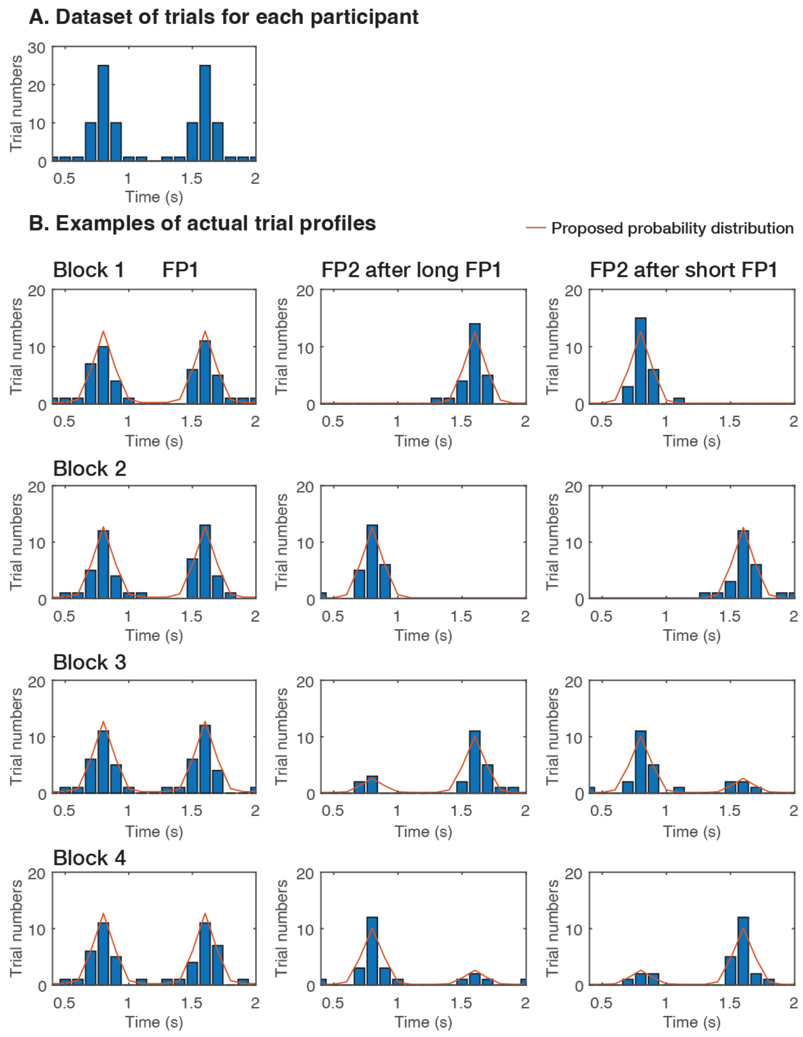


**Supplementary Figure 7 Actual trial numbers for FPs**. There were 50 trials of the two-FP sequence, resulting in a total of 100 FP trials for each block. (A) Initial trial numbers for each participant and block were determined based on a 50-50 $PDF_L$. (B) For each participant, FPs were randomly selected in the range of L and S, and paired as *LL*, *SS*, *LS*, or *SL*. The trial numbers of the four sequence types are shown in Figure 2A. An example of the actual trial numbers in one participant is shown.

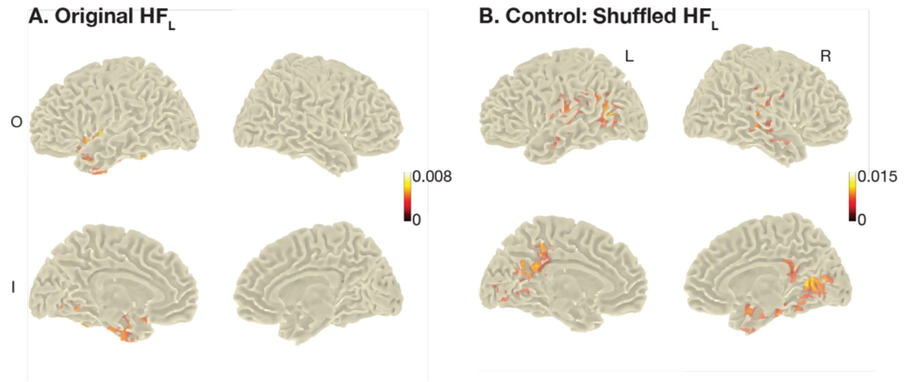


**Supplementary Figure 8 Neural correlates of original and shuffled $HF_L$.** EEG source responses during FP1 were trained against original $HF_L$ in the panel A and shuffled $HF_L$ as a control in the panel B. Four cortical surfaces with significant correlation coefficients are shown. The color bar shows the correlation coefficients.
